# Neuralace: Manufacture, Parylene-C Coating, and Mechanical Properties

**DOI:** 10.1101/2025.04.24.650335

**Authors:** Juan Pablo Botero Torres, Spencer M. Roberts, Piotr Mackowiak, Nicholas Witham, Lukas Selzer, Balaji Srikanthan, Sandeep Negi, Kai Zoschke, Florian Solzbacher

**Affiliations:** Department of Electrical and Computer Engineering, The University of Utah, Salt Lake City, UT, United States of America; Department of Materials Science and Engineering, The University of Utah, Salt Lake City, UT, United States of America; Department of Biomedical Engineering, The University of Utah, Salt Lake City, UT, United States of America; Fraunhofer Institute for Reliability and Microintegration IZM, Berlin, Germany; Blackrock Neurotech, Salt Lake City, UT, United States of America

**Author notes:** Author to whom any correspondence should be addressed.

**Keywords:** Conformable subdural implants, Flexural characterization, Neuralace, Design of experiments

## Abstract

Subdural electrode arrays have traditionally been used for epilepsy monitoring and surgical planning. With the emergence of brain-computer interface (BCI) applications, these arrays are now being explored for chronic use, leveraging their ability to record subdural signals for neural decoding. Transitioning from short-term implantation in epilepsy monitoring to long-term use in BCIs requires advancements through consideration of the foreign body response to ensure long-term durability and functionality. Biocompatibility challenges, such as fibrotic encapsulation and reactive astrogliosis, highlight the need for conformal subdural implant designs that minimize mechanical stress on neural tissue. This study investigates the mechanical properties of the Neuralace, a novel ultra-thin, high-channel-count mesh-type surface grid.

We characterized the stiffness of the silicon-based interposer electrode meshes and evaluated the effects of various geometric configurations and polymeric encapsulation layers on the mechanical performance. Using a full factorial design of experiments framework and a custom low-force four-point bending setup, we identified the design factors that impact the stiffness of Neuralace structures. The results showed highly consistent stiffness measurements. The stiffness values of Neuralace structures ranged from 2.99 N/m to 7.21 N/m, depending on the cell-wall thickness (*CWT*) of the lace, the orientation angle, and whether the structures were encapsulated with parylene-C (PPXC). Orientation and *CWT* had the largest impact on the stiffness of the structures, while the effects of PPXC encapsulation were statistically significant but more subtle.

The stiffest Neuralace configuration is expected to exert forces approximately 10 to 100 times lower than commercially available subdural implants would when conforming to radius of the smallest anatomical gyrus. This work demonstrates the feasibility of tailoring the mechanical properties of Neuralace to improve its suitability for chronic neural implantation, providing insights for future design iterations and conformable implant development.

## Introduction

Subdural electrode arrays come in a variety of form factors. Clinically used Electrocortical Grid (ECoG) and Micro Grid (μECoG) arrays have been used for decades to record electrical brain activity from the cortical surface in the form of local field potentials (LFPs). Subdural arrays are used in epilepsy monitoring to identify and localize seizure foci and to inform surgical resection areas for refractory/anti-epileptic drug (AED) treatment resistant patients. In scientific research publications, typically, thin film based micro surface grids have been used to record LFPs and single unit action potentials e.g. for better seizure foci location using high frequence oscillations or increasingly for the use in controlling external devices as part of a brain computer interface (BCI).

In BCIs neural activity recorded by implanted electrode arrays can be decoded into control signals that allow patients with severe motor impairment (i.e., amyotrophic lateral sclerosis, tetraplegia, and spinal cord injury among others) to control a computer^1–3^, control robotic prosthetics^4^, and even bridge spinal cord injuries^5^. Epilepsy monitoring is currently a sub-chronic application in which the implant remains in the patient for less than 30 days. Future use cases in epilepsy monitoring where the recorded signals are used to drive a neuromodulation treatment strategy, as well as BCI applications require chronic implantation, with the implant ideally remaining in the patient for decades. Therefore, to be relevant to BCIs, subdural electrodes must be designed to have vastly longer implantation lifetimes than what is currently available.

A key factor determining the lifetime of neural implants is their biocompatibility. *D*.*F. Williams* defines biocompatibility as the capacity of a device to adequately perform with an appropriate host response in a given application^6^. For subdural implants, the appropriate host response is the local foreign body response (FBR) triggered by the device. It has been shown that following intracortical electrode implantation, astrocytes and microglia are activated, inducing a morphological transformation into their reactive type, characterized by an increase in the cell volume, cell count, and their inflammatory reactions^7^. Chronically, such reactive cell types aggregate in the electrode-brain tissue interface, resulting in a glial scar that not only isolates the active electrode sites from the signal source, but may also lead to dendritic retraction, neuronal death, and creates a harsh acidic and oxidative environment which can corrode electrode sites and degrade polymeric encapsulations^8,9,10^. Such degradation can reduce the ability of intracortical arrays to record neural signals by increasing the impedance of the electrode-brain tissue interface. This chronic response is often referred to as reactive astrogliosis that leads to glial scarification, and it has been shown to be the prevalent FBR for intracortical electrodes.

However, for subdural implants extensive literature searches have only identified six studies that performed immunohistological assays to evaluate the *in vivo* chronic FBR ^11–16^. The limited number is likely due to complexity, cost, and limited clinically relevant chronic use cases. Nonetheless, these studies show that subdural implants will undergo fibrotic encapsulation (FE) upon implantation, which results from the aggregation of multinucleated foreign body giant cells and mononuclear macrophages. It is important to note that FE develops independently from glial scarring (i.e., using different cellular mechanisms and cell types); however, how these types of FBR interact is unknown. *Ryapalova et al. and Schnedel et al*. report that only FE is present after implanting a Rhesus macaque and four rats for two years and one year, respectively. While *Henle et al*. and *Degenhart et al*. report FE and astroglial remodeling (i.e., increased density of microglia and/or astrocytes in the vicinity of the implant) after implanting five rats and a Rhesus macaque for 25 and 95 weeks respectively. And *Yan et al*. have performed two studies in which they report FE and signs of reactive astrogliosis (i.e., astroglial remodeling and signs of reactive activation) after implanting six beagles and two monkeys for six and eight months, respectively. It is important to note that the astroglial remodeling and signs of reactive astrogliosis reported in these studies is only present in the brain tissue underneath the subdural implants.

Overall given the complexity and confounding factors involved with *in vivo* evaluation (i.e., animal model, surgical procedure, implantation time, implant materials, implant design) and due to the lack of standardization on the parameters reported in these studies, the outcomes on whether reactive astrogliosis is present and to what extent are non-conclusive. However, *Moshayedi et al*. showed that microglia and astrocytes are mechanosensitive cells, which can be activated into their reactive state due to changes in the mechanical load imposed by their surrounding environment^17^. Thus, there is a physiological pathway for chronically implanted subdural electrodes to trigger reactive astrogliosis via mechanical stress.

Subdural neural implants are subjected to different mechanical loads throughout their lifetime. The first loads that neural implants experience are those exerted during surgical manipulation, which are considered to be the highest mechanical stress that the implant must endure. Once the implant is in place, it must conform to the geometry of the brain, stressing the composite material; resulting in a non-uniform load that varies depending on the size of the implant and the region of implantation. During chronic use, the implant is also subjected to cyclical loading induced by respiratory and vascular cycles that produce the so-called brain micromotions. It has been shown that in adult rats, these micromotions can generate up to 30 μm displacement at the probe-tissue interface^18^. In humans, brain motion analysis has been performed using different MRI analysis techniques, showing a displacement of up to 500 μm varying by brain region^19,20^. Overall, neural implants must resist the mechanical stress caused by implantation, conformation to the brain topology, and micromotions.

The flexural stress endured by the implant leads to mechanical loads exerted on the surface of the brain cortex. Which, as described earlier, may trigger reactive astrogliosis. Furthermore, it has been shown that exerting force on neural tissue is correlated with decreased action potentials^21^, can cause reduced conductivity on neurons^22^, and even lead to neuronal death^23^. Thus, it is hypothesized that reducing the mechanical strain on the brain tissue could reduce potential adverse effects. This has led researchers, focused on neural implant development, in a pursuit to reduce such loads. Several approaches have been reported in literature, mainly utilizing materials whose Young’s modulus is closer to neural tissue (e.g., polymers, hydrogels, etc.), and/or reducing the thickness and footprint of the substrate, such as to reduce the effective forces acting on the neural tissue. Reviewing the state-of-the-art of conformable neural implants falls out of the scope of this work, but there have been several in-depth reviews published that the reader can refer to ^24,25^.

This work characterizes the mechanical properties of a novel device architecture, the Neuralace device (Blackrock Neurotech) and the impact of various geometric configurations and varying encapsulation coating layers as key component driving chronic bio-compatibility of the device. The Neuralace is currently a silicon or polymer based, ultra-thin (10-40 μm thick) and ultra-high channel count (>10,000 electrodes) mesh-type surface grid with integrated electronics for signal processing and wireless data transfer. The mechanical characterization was performed on the Si based interposer and does not consider any active layers or components (i.e., metallization layers and doped regions), as these layers do not contribute significantly to the bulk volume of the structures. Thus, the anticipated impact of these layers on the mechanical properties of the structures is minimal.

At this stage of development, the design space is large and can be difficult to explore efficiently. Typically, design optimization can be explored via experimentation or modeling. Computational models, such as finite element analysis (FEA) models, could be used to explore a narrow optimization space. For example, they could be used to cheaply explore the effect of different lace patterns on the stiffness, or the effect of different coatings. However, it can be difficult to accurately produce FEA models of devices with multiple material boundaries, thin film defects, large deflections, and extreme aspect ratios, all of which can apply to the Neuralace structure^26–29^. On the other hand, experimentation can be costly and provide limited or noisy information if not well designed. Design of experiment (DOE) is a statistical method that accounts for resource limitations and anticipated analysis techniques. Typical DOE methods include, response surface methodology (RMS), space filling designs, and Taguchi arrays, among others. Nevertheless, all DOE methods follow the same step-by-step framework: describe your goal and identify responses of interest and potentially impactful factors, specify an assumed model of the physical situation, generate a design that is consistent with the assumed model, collect data with the trial settings specified by the design, fit your assumed model to the experimental data, and finally use the refined model to address the experimental goals, such as identifying the factors of impact and their optimal levels. We follow this methodology in this work by using a full factorial design to explore the design factors of the Neuralace that have the largest impact on structure stiffness. Our goal is two-fold: first, to narrow down the stiffness optimization space and identify the range of levels that should be considered with subsequent Neuralace iterations. Second, we aim to characterize the stiffness of Neuralace structures and evaluate its mechanical performance relative to other conformable brain surface implants.

To perform the mechanical tests, we developed a custom low force four-point bending setup as no off-the-shelf instrumentation is designed for measuring the small forces (μN to mN) associated with imposing flexural loads on conformable neural implants. On the following sections, we will show the microfabrication process of Neuralace structures and their mechanical characterization, focusing on the effect of the polymeric encapsulation (i.e, PPXC), the orientation of the bending load, and the width of the traces on the mesh-like hexagonal structure.

## Methodology

### Fabrication of Neuralace structures

Figure 2 shows the simplified Neuralace manufacturing process. First, the SOI wafer (1 μm buried oxide (BOX) and 10 μm thick device layers) of is bonded face down to silicon carrier wafer using an adhesive (Pieplow & Brand, Kunstharzklebekit MKS). The carrier wafer of the SOI wafer is then thinned by means of wafer grinding. This is a mechanical process that introduces high stress into the wafer surface, so this process is stopped approximately 30 μm before the buried oxide (BOX) layer is reached. The remaining silicon is removed via reactive ion etching (RIE) with a SF_6_ flow of 533 sccm in a Dual Source System with powers of 3000 W and 2000 W. Bias is set to 0 W and the chuck is maintained at 20°C with 15 sccm of He flow for backside cooling. The BOX is then removed by RIE as well, with C_4_F_8_ and He flows of 15 and 174 sccm, respectively.

Chuck temperature is maintained at 0 °C and the bias is 400 W. ICP power is set to 2000 W. The now exposed device silicon layer is coated with Microchemical AZ 10XT photoresist and the lace structures are transferred to the resist by means of UV exposure and a photomask. Subsequently, the lace pattern is etched into the silicon via deep reactive ion etching (DRIE), where the resist protects against the etching attack, while the open areas are completely etched, and the structures are thus transferred to the silicon. DRIE is performed via a three part etch process, which includes: a C_4_F_8_ deposition step, an towards the directed O_2_ etch, and an SF_6_ etch. For the C_4_F_8_ step, chamber pressure is 45 mTorr and flow rates are 400 sccm + 100 sccm (dual feed system). ICP power is set with the Dual Source System at 2500 W and 500 W and the bias is set to 0 W. The chuck temperature is maintained at 20°C. For the O_2_ etch, the chamber pressure is 28 mTorr and the bias is 400 W. All other parameters are the same as the C_4_F_8_ deposition step. For the SF_6_ etch step, the chamber pressure is 80 mTorr and the gas flow rates are 500 sccm + 1 sccm (dual feed system). The bias is set to 50 W. The ICP power and chuck temperature are the same as the previous two steps. These steps are repeated for 28 cycles. The process times within one cycle for the three steps of deposition, first etch and second etch steps of the etching process are 1.0, 1.3 and 2.0 seconds.

After the DRIE, the photoresist is removed with an O_2_ RIE plasma, and the devices are released from the carrier wafer via wet stripping with Microchemicals AZ100 (1:1 ratio with DI water) for 7 minutes at 50° C. After the solvent bath, the samples are rinsed in DI water and dried with nitrogen. To remove residual adhesive, the individual lace structures are placed in a perforated Teflon carrier for a 30 minute soak in a 120°C (H_2_SO_4_/H_2_O_2_) Piranha solution bath. Following the Piranha solution bath, the laces are rinsed with a DI water. The laces are collected out of the carrier one-by-one with tweezers and dried on a cleanroom tissue under laminar nitrogen flow.

Before any bend testing, a thermal oxide is grown on the laces in a 1000 °C thermal oxidation furnace. The structures are clamped between silicon slices wafers which are positioned horizontally on a glass carrier and placed in the furnace. White light spectrometry is used to characterize the oxide thickness. Finally, after initial bend testing, the structures are coated with PPXC via chemical vapor deposition (CVD). To conformally coat the devices with PPXC, structures were suspended on four contact points in custom 3D holder that enables the flow of vaporized dimer above and below the structure. The design of the holder minimizes non-uniformity by minimizing contact area with the structures. Microscopy imaging of the coated structures showed that the diameter of contact area was 216 ± 69 μm across all coated structures (see S2). After deposition, the thickness of the PPXC layer was characterized on Si witness chips that were coated alongside the Neuralace structures. This characterization was performed with a profilometer (Dektak Xt stylus, Bruker) and resulted in an average thickness of 1.78 μm ± 20 nm (see S2).

### Design of Experiments

Quality performance studies conducted by medical device manufacturers typically follow six sigma methodologies, which seek to limit defects to 3.4 per million samples/tests^29^. Characterization studies with those confidence levels require much larger sample counts than would be possible or warranted in an academic setting. Nevertheless, sufficiently powered (80 % or more) effect screening studies are vital to identify which experimental factors impact your desired device response and what effect size(s) should be expected. We designed a full factorial experiment to screen for the effects of lace structure *CWT* (25, 50, 75, and 100 μm), structure orientation (0° and 90°), and PPXC thickness (0 and 1 μm) on the stiffness of the Neuralace device, determined by bend testing. The full factorial combination of these levels produces 16 (4 × 2 × 2) unique experimental conditions. Three samples were used per *CWT*, leading to a total number of 12 individual samples for this study. Samples were first measured without PPXC coating and then with PPXC coating. Each unique bending trail was repeated three times by resetting the sample to increase statistical power, leading to a total of 144 (16 × 3 × 3) individual bending trails. Four samples, one for each *CWT*, were randomly pooled together as one batch, leading to three batches (E1, E2, E3), to examine the repeatability of the experiment and sample ID influence. In this study, each of the unique 16 experimental conditions was tested 9 times, giving the study a power of 85 % with a significance level (α) of 0.05 and a difference to detect of one standard deviation ().

### Custom mechanical testing setup: design and validation

The flexural bending forces of the ultrathin Neuralace structures are too low for most commercially available bend testing systems, so we designed and evaluated a custom low-force, four-point bending setup to measure their flexural properties (see Figure 3). The components of the system include: a precision scale with a resolution of 9.8 μN (VWR 160AC) as the force transducer, an automated z-stage micromanipulator (Thorlabs PT1-Z9) with an incremental resolution of 0.2 μm, a manual x-y stage (Thorlabs ST1XY-S), a displacement sensor (AR700-0125, Acuity Laser) with a linearity of 0.95 μm, and custom 3D printed loading geometries. The z-stage is mounted in the x-y stage and secured to a vertical optical breadboard via a 3D printed (CF infused PLA) L-shaped bracket. The geometry of the loading and supporting noses critically impact the results of a bending experiment. We used four-point bending so the maximum stress is uniformly distributed between the loading noses. ASTM standards recommend that the support span should be long enough to allow for an overhang of at least 10 % of the structure length on each end^30,31^. Given that the Neuralace structures are squared with 11 mm sides, we choose a gap span of 9 mm. Loading spans of both one-third and one-half of the support span have been widely used in literature^32–34^, and are accepted by international standards^30,31^. However, we opted for one-third loading spans to increase sensitivity, as the load is distributed closer to the center of the structure, inducing higher stress at lower strain. Finally, our system uses metallic hypodermic needles as curved loading and spanning surfaces to reduce friction and possible surface cracks to the test structure. Experimental runs and data acquisition are automated with Python. The software requests the operator to input experimental hyperparameters, performs semi-automated alignment, records raw and steady-state force and displacement values, and displays data in real-time for monitoring purposes. The four-point bending setup was validated by testing solid Si beams with the same dimensions as the Neuralace structures (11 mm x 11 mm x 10 μm), calculating the stiffnesses and converting them into Youngs Modulus (*E*) accounting for the geometry as detailed in ^34^. This resulted in *E* = (166 ± 5) GPa (N=9), which aligns with the literature reported value of 169 GPa. For those who which to reproduce the testing setup the code, schematics, and validation data for the system can be obtained upon reasonable request to the authors.

### Flexural characterization: experimental procedure

Flexural characterization was conducted non-destructively in a physiologically relevant displacement range. In previous work, we defined the maximum bending stress in a physiological environment—ignoring acceleration-induced deformations—by evaluating the radius of curvature of the different gyri of the brain cortex in the MRI scans of a Caucasian 38-year-old female^34^. The maximum stress induced by conformation of the subdural implant to the gyrus with the smallest radius, found to be 60 mm. Considering the dimensions of the Neuralace, this stress is induced at approximately 160 μm of displacement, which is within the elastic deformation region of these structures.

Prior to imposing bending loads on the test structure, the alignment of the loading geometries was verified using a 3D-printed plate with parallel grooves, see section one of the supplementary material (S1). After alignment adjustments were completed, the software recorded the initial position of the z-stage, instructed the operator to remove the alignment tool, and tared the scale. The structures were then placed on the lower loading geometry, and the upper loading geometry was displaced in steps of 10 μm (the smallest displacement allowed by the micromanipulator) until contact was detected, defined as when the force exceeded 100 μN (~10x the resolution of the scale). Once the experimental contact point was established, data acquisition began, and displacement advanced in 20 μm steps to the max of 160 μm with 10 second holds after each step to allow perturbations to dissipate. The raw data of force and displacement were recorded continuously and steady-state data was averaged during each 10 seconds hold. Each structure was subjected to five loading and unloading cycles during a single bending trial.

### Statistical analysis

To analyze the full factorial experiment, we used a standard least squares linear regression model to conduct an effects screening, which examines the impact of each factor and their second order interactions (i.e., predictors) on the stiffness response. Statistical data analysis was conducted with JMP Pro 17 (JMP). To briefly summarize the algorithm JMP uses, the model is parameterized with an *n* by *p* design matrix *X* where *n* is the number of observations and *p* is the number of predictors. The linear model is described in matrix notation in the following equations, where *β* is the vector of regression coefficients, 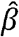 is the estimate of that vector, and *Y* is the vector of observed responses (i.e., measured stiffnesses).

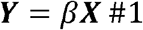

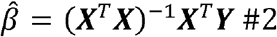

For an effect screening analysis, the estimate vector is used to evaluate the potential impact of each predictor. The effect of a predictor is assumed to be zero, the estimate vector is modified accordingly, and a modified estimated response vector is calculated. The error of the modified estimated response vector is compared via *t*-test to the unmodified response vector and a *p*-value ≤ 0.01 indicates the predictor has a statistically significant impact on the response. Because multiple predictors were tested, we used the false discovery rate (FDR) technique (α= 0.01) which gives more conservative estimate of the statistically significant impact of each predictor^35^.

## Results

### Flexural characterization of Neuralace

To characterize the flexural properties of Neuralace structures we performed cyclical four-point flexural bend tests. The force deflection curves of each bending trial were fitted with an ordinary least squares regression to find the stiffness, as shown in Figure 4A. Within a bending trial the stiffnesses of each loading and unloading cycle (5 per trial) are tightly grouped as illustrated by the grouping of the colored markers in Figure 4B. This spread within bending trials was quantified by calculating the coefficient of variability (CV), defined as the standard deviation divided by the mean. The intra-trial CV was calculated using the stiffness for each loading and unloading cycle within a bending trial. This resulted in a mean CV of 0.57 % (σ = 0.53 %), which validates the stability of the cyclical loading experimental procedure. Because the cycle stiffnesses of each trial are so tightly grouped, the data from all the cycles were used for the least squares fit of a single trial and used to analyze the overall impact of each experimental factor on the mechanical properties of the structures. Across repeated bending trials of any given experimental condition, the stiffnesses are randomly distributed within structures, as seen by the representative violin plots in Figure 4B. Additional violin plots can be found in the supplemental information (S3). This inter-trial variability was also quantified with CV, which was calculated for each experimental condition across repeated bending trials. The mean CV across all conditions was 5.15 % (σ = 3.15 %), ranging from 0.38 % to 14 %. This indicates that on average the repeatability across repeated bending trials is adequate. However, the CV analysis exposes a random variability on repeated trials that is due to manual sample placement and alignment. The variability between repeated placement was evaluated post-hoc by imaging ten repeated sample placements per target orientation (0° and 90°) and calculating the difference between the mean orientation and the sample orientation (see S2 for more details). The analysis suggests that the orientation angle was controlled with a resolution of 3°. Figure 4C shows the aggregated data per experimental condition, where each box plot consists of 9 stiffness across 3 trials of 3 structures. The red and blue colors indicate 1 μm of PPXC or no PPXC coating, respectively. The darker and lighter saturations indicate a structure placement of 0° or 90°, respectively. There is a general upward trend in stiffness as the lace *CWT* increases and a large difference in stiffness between a 0° and 90° orientation. However, the effect of PPXC encapsulation is subtle increase on stiffness. The mean stiffness measured for the Neuralace structures with the same experimental conditions ranged from 2.99 N/m (*CWT* = 25 μm, orientation = 90°, uncoated) to 7.21 N/m (*CWT* = 100 μm, orientation = 0°, PPXC coated). Mean stiffnesses for all experimental conditions and other descriptive statistics summarize the obtained results (see Table 1).

**Table 1:**
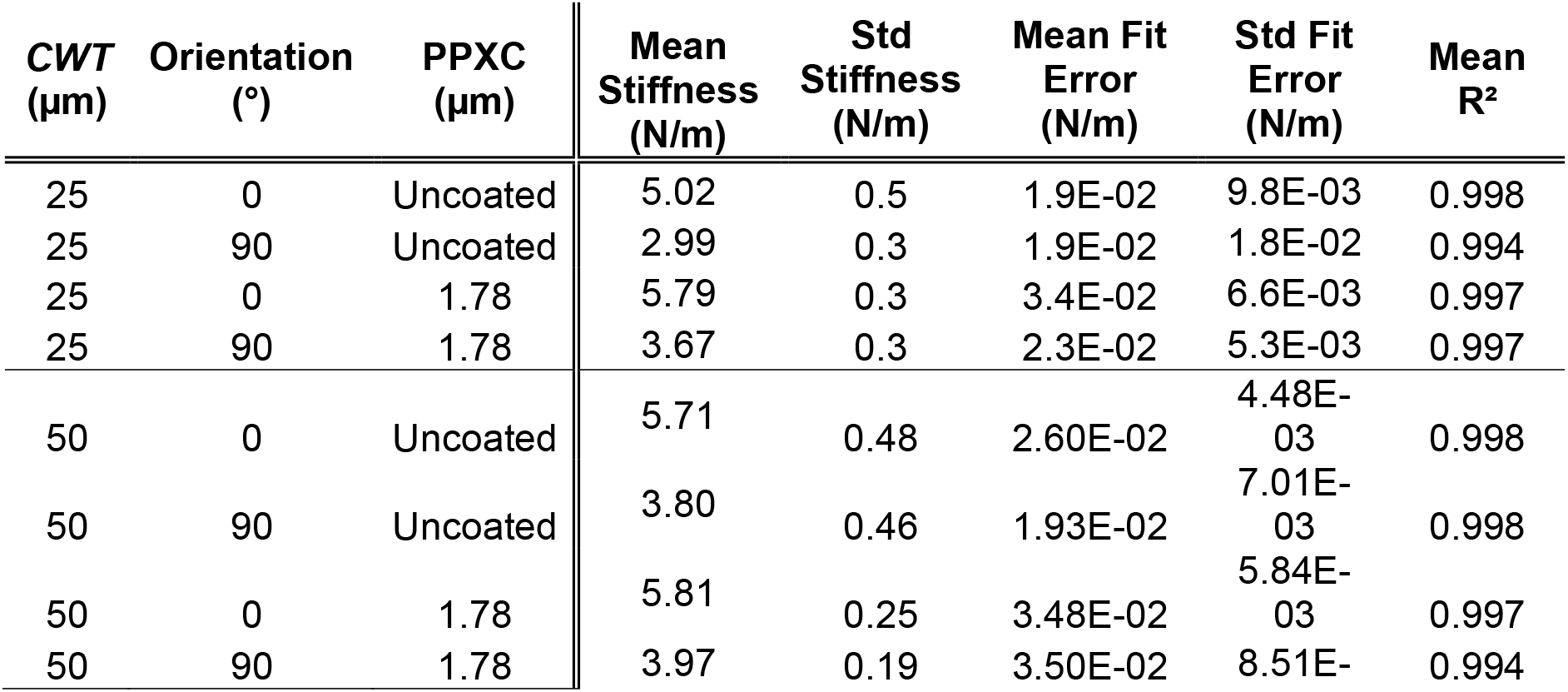
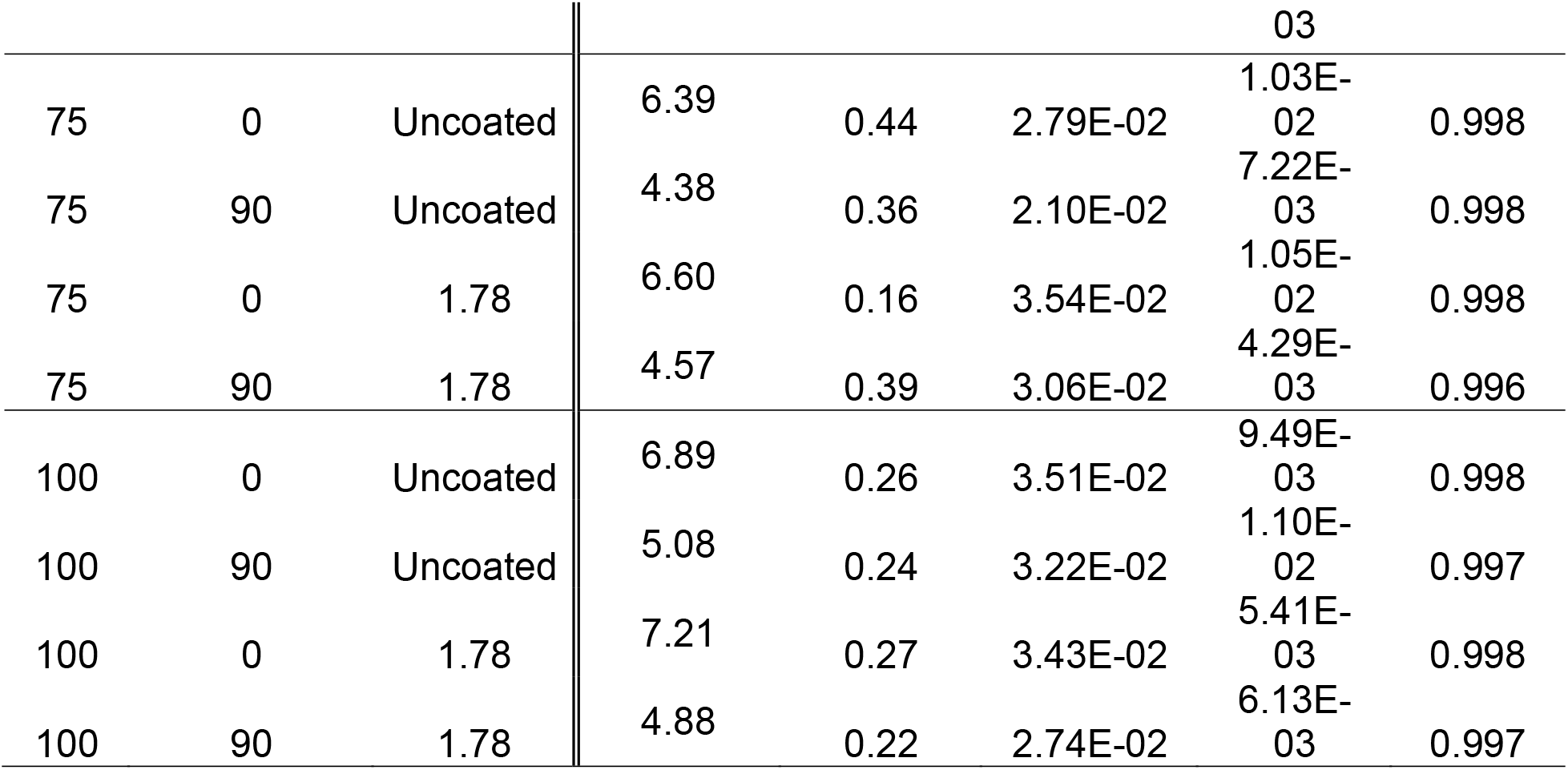
Stiffness fits and statistics (N=9) for each 4-point bending experimental condition (i.e., row)

### Mechanical performance comparison to other subdural implants

For comparison, the maximum forces expected upon implantation of other sub-dural implants was taken from previous work and plotted alongside the forces of the stiffest Neuralace configuration (*CWT* = 100 μm), see Figure 5^34^. The comparison maximum forces shown were also estimated based on the conformation to a 60 mm gyrus, as detailed in the flexural characterization section of the methodology. The forces exerted by the Neuralace structures are about 100 times less than the commercially available devices tested and about 10 times less than liquid crystal polymer (LCP) films. For context, the forces exerted by these Neuralace structures are approximately two time less that those that would be exerted by a solid Si beam of the same dimensions.

### Effects screening

To further inspect the impact of the experimental factors on the stiffness of the Neuralace structures, an effect screening study was performed, as detailed in the DOE section of the methodology. A visual inspection of Figure 4C does not elucidate a clear effect of PPXC on stiffness, but does suggest a slightly positive trend. The effects screening summarized in Figure 6 provide more illumination of this phenomenon. The FDR Logworth in Figure 6A shows the relative effect strength of each factor and one second order interaction (PPXC thickness \**CWT*). The vertical blue line indicates an FDR logworth value 2, which corresponds to a statistically significant FDR *p*-value of 0.01. Therefore, out of the four primary factors, only the experiment ID factor did not have a significant impact on stiffness. Orientation and *CWT* had the largest impact on stiffness by far, though PPXC thickness and a second order interaction between PPXC and *CWT* were also significant but by a very reduced margin. This can be visualized in Figure 6C by the change in edge shape along the *CWT* axis. The two surface plots in Figure 6B and 6C show stiffness response surfaces for the most significant factors.

## Discussion

### Using effects screening as a design guide

The purpose of the effects screening facilitated by JMP Pro is to make further DOE more efficient and provide a window into the optimization space covered by the factor levels. Within that window, the predictive accuracy of the linear regression model developed by JMP is limited by the two-level factors, such as orientation, which physically will have a sinusoidal impact on stiffness instead of a linear impact. Nevertheless, the model remains useful in providing feedback on the study and guiding iterative design choices. For example, only experiment ID factor did not have a significant impact on stiffness, suggesting that our system and fabrication protocol are well controlled and reproducible. On the other hand, the effects screening indicate that orientation and *CWT* are the most important factors to control in subsequent studies. Furthermore, the model elucidates a small but statistically significant second order interaction between PPXC thickness and *CWT*. These clues suggest the next iteration of Neuralace designs should consider— among other things—the following questions: what is optimal orientation of the device to the curvature of the brain, should the device be symmetrical to simplify the implant procedure, whether a reduction in *CWT* below 25 μm is worth the reduction in stiffness, should the *CWT* be increased so that subsequent layers have less effect on the stiffness, what thickness of PPXC can be deposited before having a larger effect on the stiffness, and which factors can be eliminated from further study and which factors merit more levels. Proper DOE and effects screening provide statistical foundations for such experimental insights.

### Anisotropy of Neuralace Structures

Orientation is the most impactful factor on the stiffness of these Neuralace structures. The anisotropy observed could arise from several factors, such as the intrinsic material properties, the mechanical properties of regular hexagonal cellular solids, and the geometry of the structures. Si is an inherently anisotropic material; the crystal structures for 100 Si wafers (as those used to fabricate these Neuralace structures) have two axes of symmetry. Therefore, there is no expected anisotropy under 90° rotations. The mechanical properties of regular hexagonal cellular solids are generally isotropic and have been thoroughly described by *Gibson et al* ^36^. However, their isotropic behavior depends on the mechanical properties of the bulk material, thus there could be an interplay between the intrinsic anisotropy of Si and the properties of the hexagonal cellular solid. Producing an analytical description of the compound effect of the cellular geometry and the crystal structure of the substrate would be a complex and time-consuming endeavor; hence, the empirical approach conducted in this study. Nevertheless, we anticipate that the main source of anisotropy is simply the non-symmetrical distribution of the cellular and solid regions. In the 0° orientation the load bearing area is dominated by the wider portion of the frame, while in the 90° orientation is dominated by the narrower portion (see Figure 1B).

**Figure 1.**
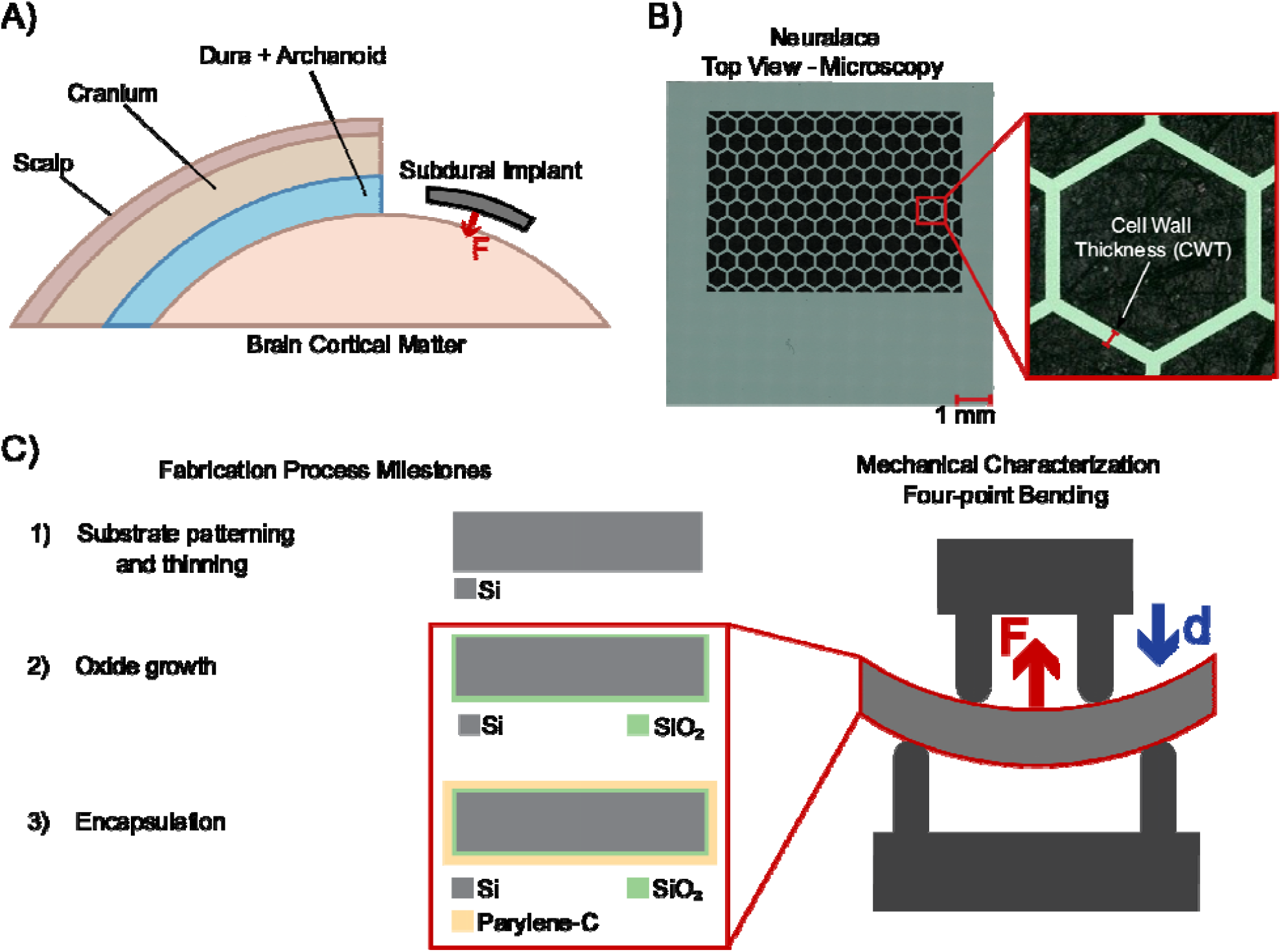
A) Schematic of the forces exerted by the Neuralace upon subdural implantation. B) Microscopy images of Neuralace structures with zoomed in region to one of the regular hexagonal cells that form the lace region. C) Cross sectional view of Neuralace structures along the fabrication milestones, red bounding box demarks the composite that were tested using four-point bending.

**Figure 2.**
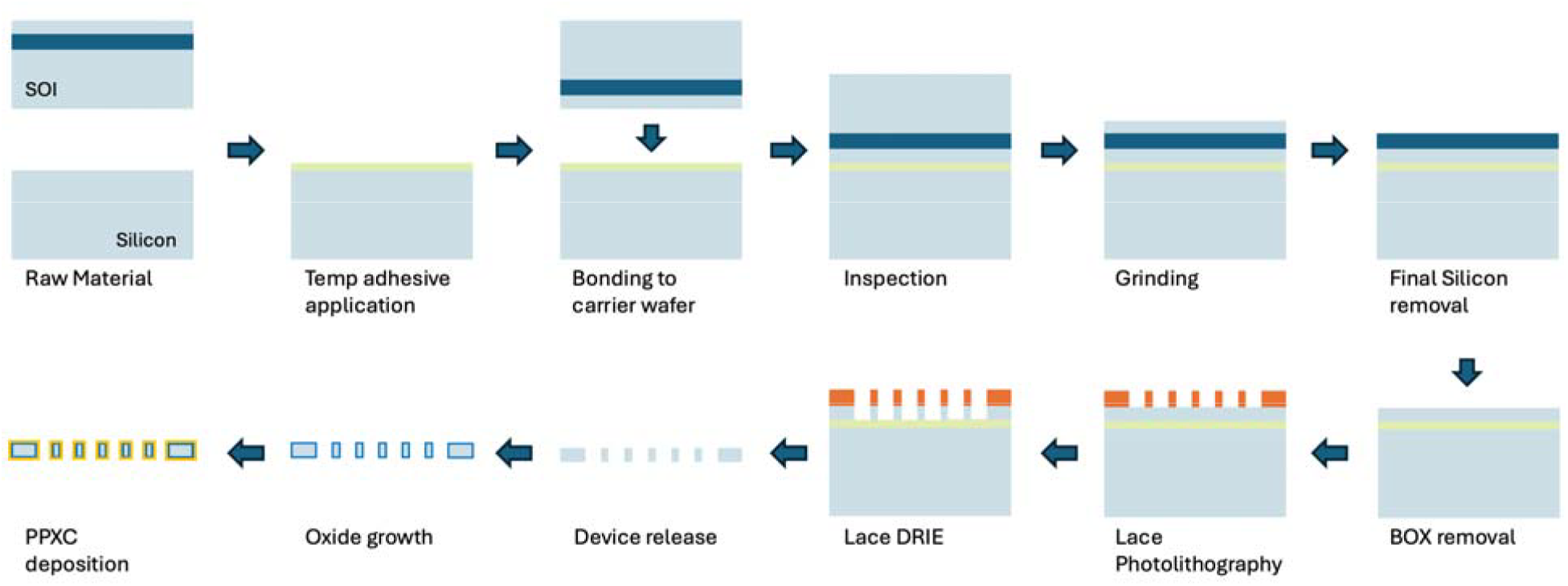
Overview of the Neuralace structure fabrication process. The final removal of the silicon layer and the BOX are achieved with RIE. The etch of the lace structures is achieved with an DRIE Bosch process. Following the structure release, a SiO_2_ layer is deposited. After initial testing, a PPXC layer is deposited via CVD (see S2).

**Figure 3.**
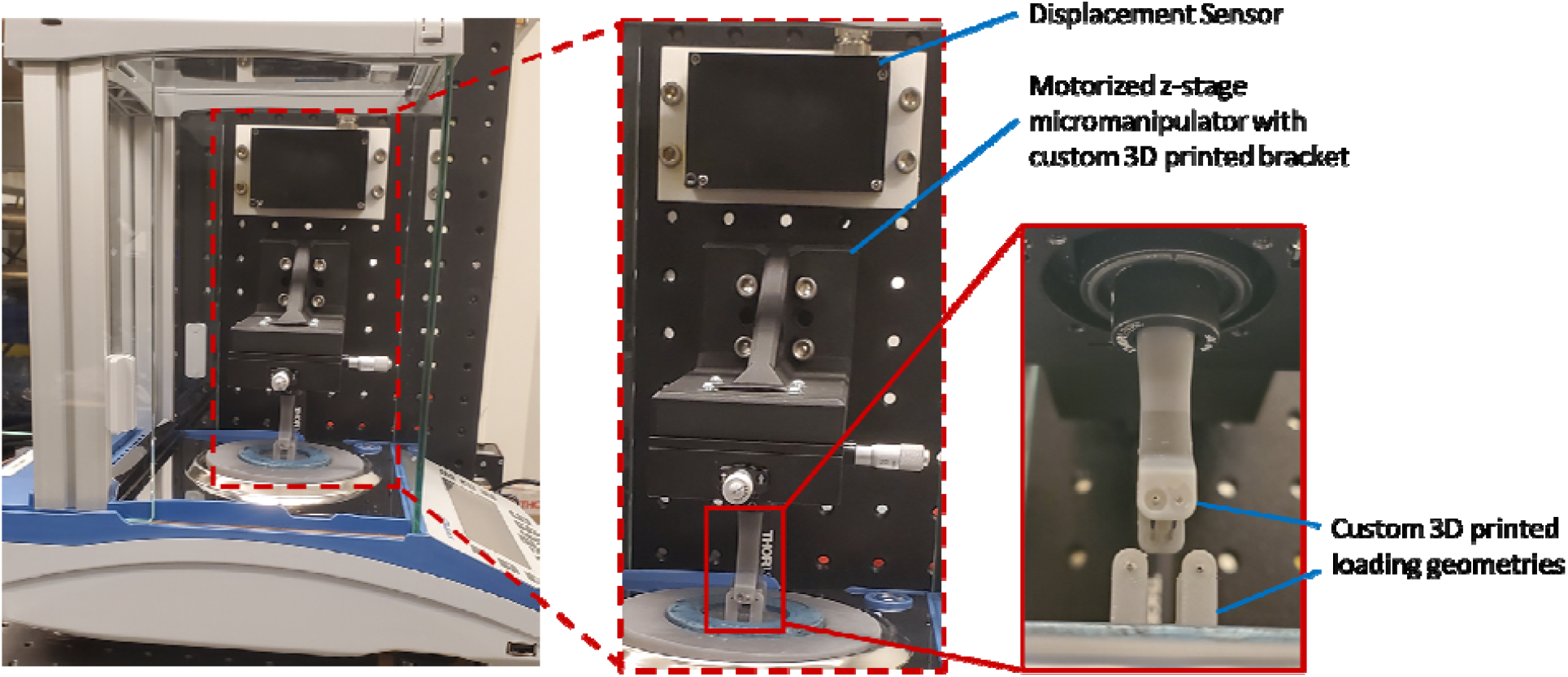
Custom flexural testing setup designed for measuring the low forces (hundreds of μN to mN) associate with the deformation of conformable subdural implants.

**Figure 4.**
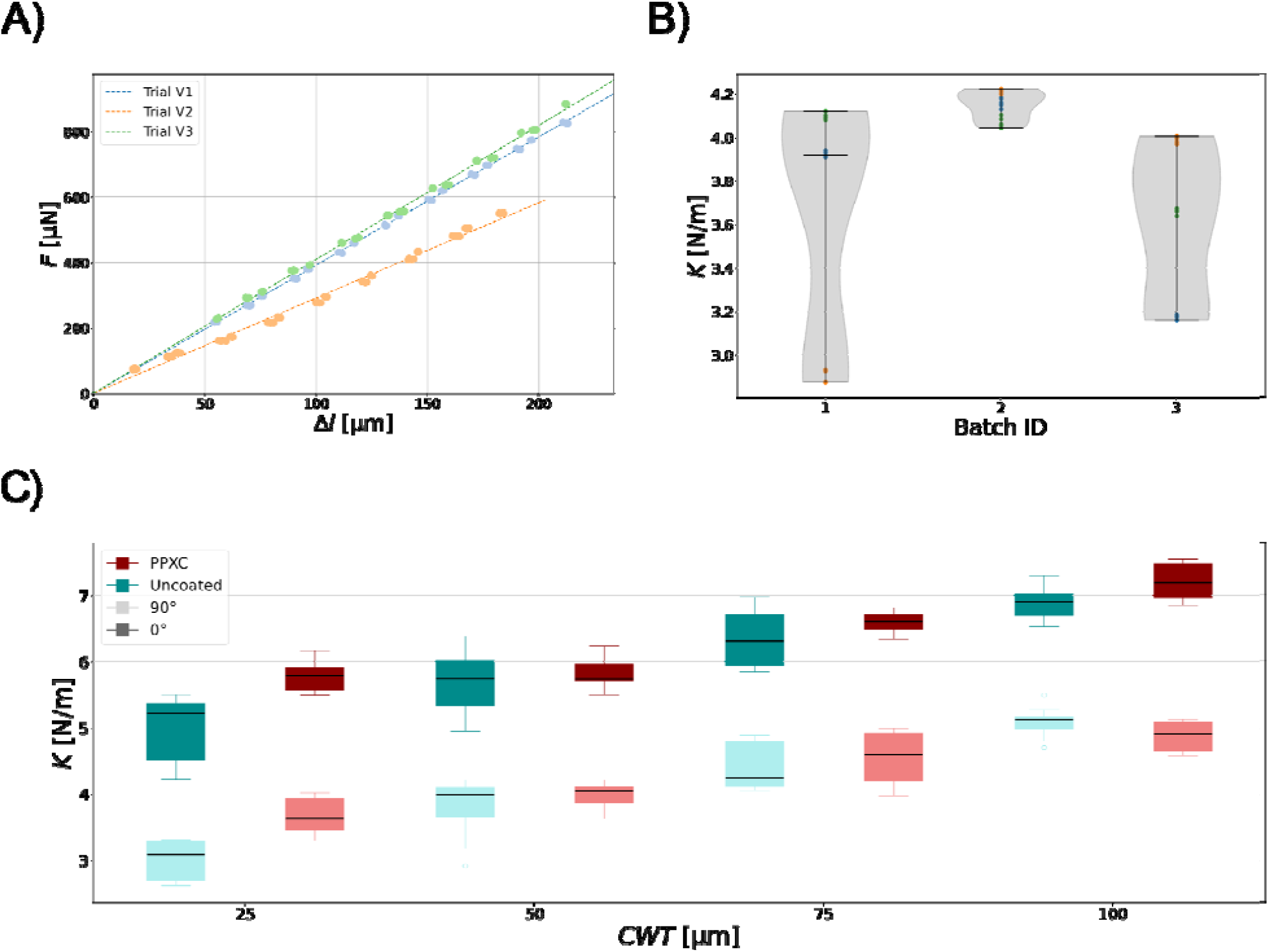
Dataset Structure A) Force-displacement curves for the three bending trials with the same experimental conditions (*CWT*=50 μm, orientation=90°, uncoated, Experiment ID= 1), the dotted lines show the linear regressions of each trial considering data from all loading and unloading cycles. B) Distribution of stiffnesses obtained from each loading and unloading cycle within an experimental trial experimental repeats of the condition (*CWT*= 50 μm, orientation=90°, uncoated)). C) Stiffness comparison between all experimental conditions.

**Figure 5.**
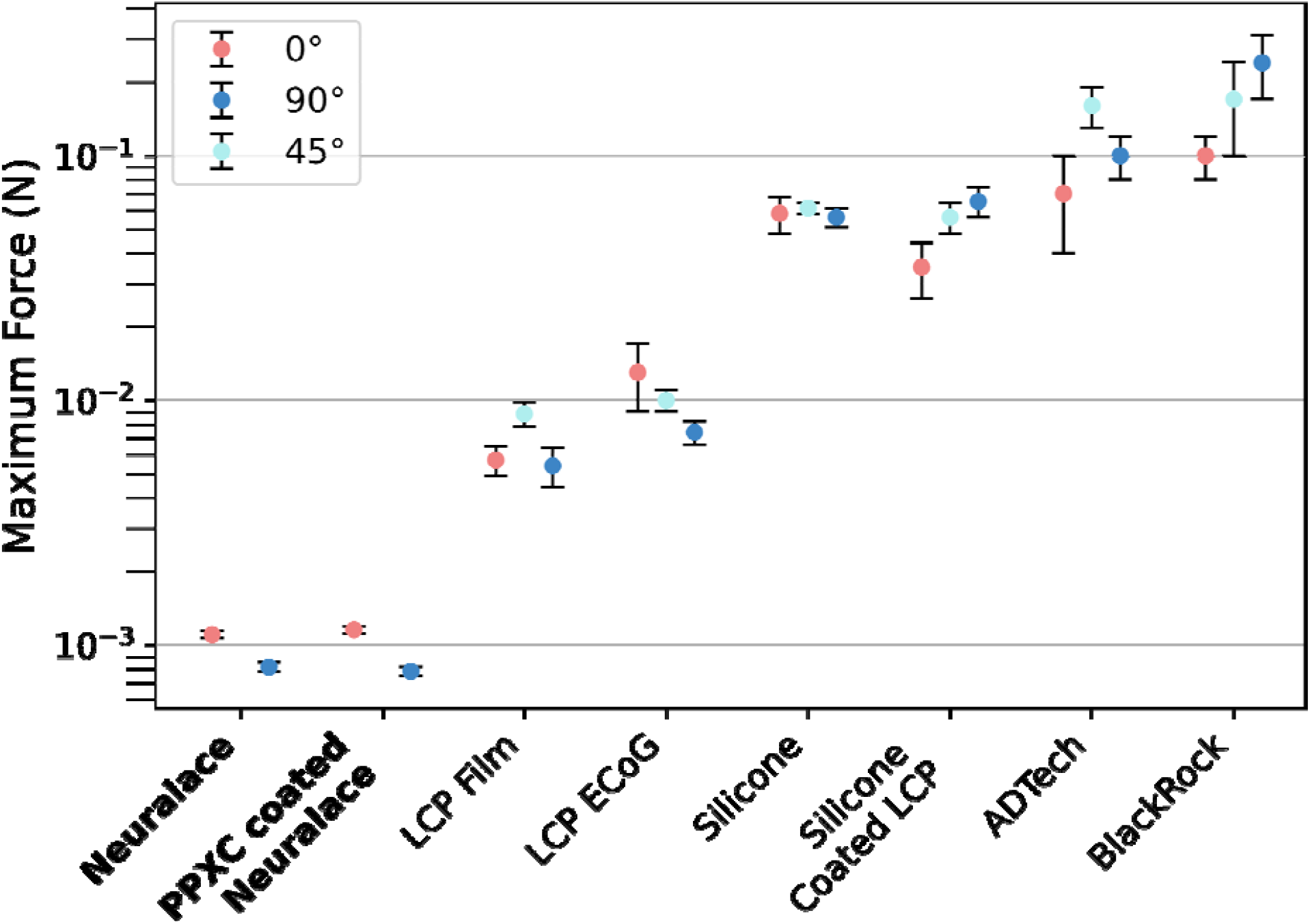
Maximum expected forces exerted by commercially available and research-grade subdural implant whe conforming to a 60 mm gyrus. Bold marks denote the data obtained in this study, all other data points where reporte by Witham et al. ^*34*^

**Figure 6.**
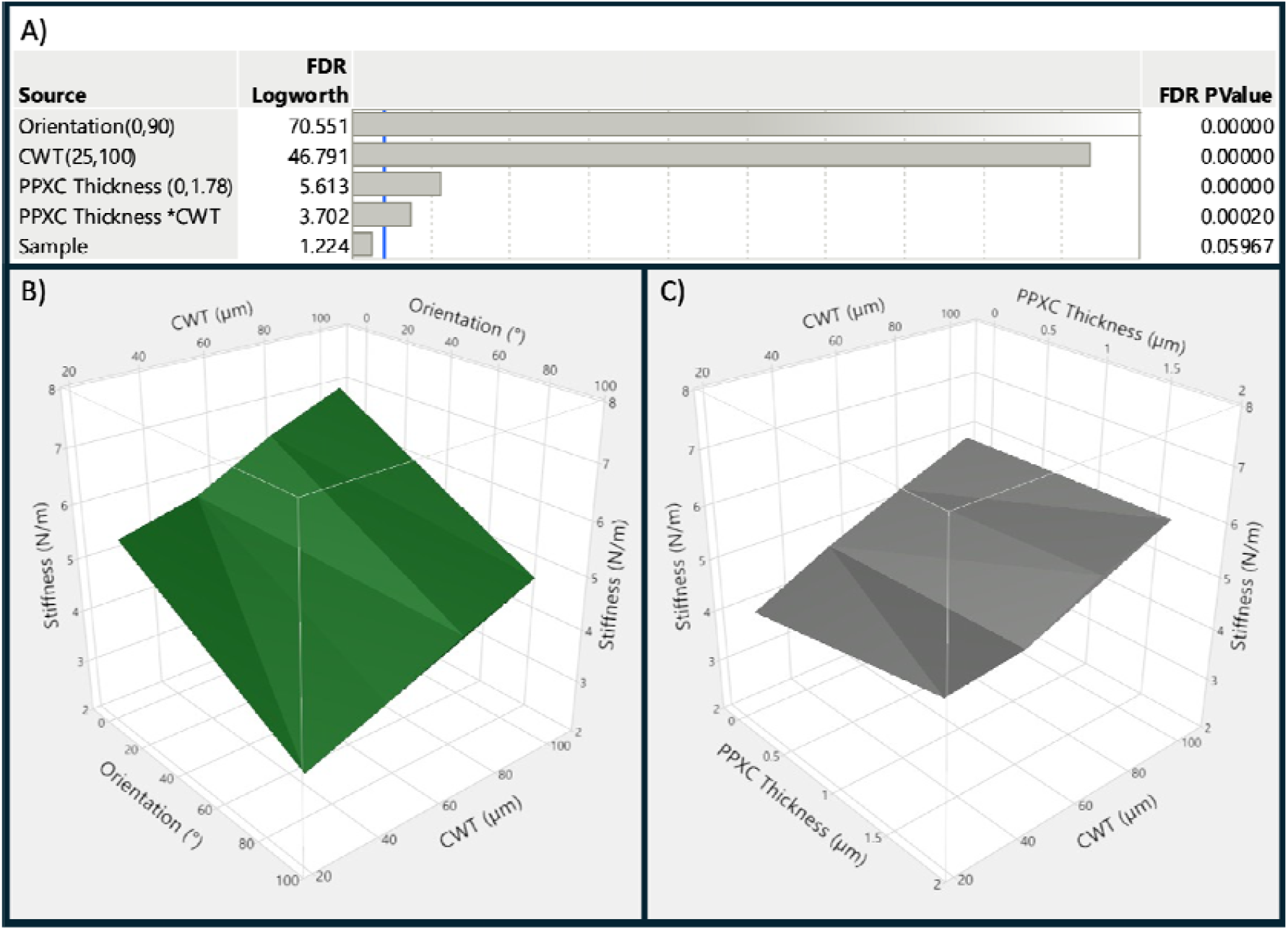
Effects screening summary and surface response plots. A) The effects summary of the various factors an second order interactions on stiffness. A false detection rate (FDR) logworth > 2 is statistically significant. Only statistically significant and primary factors are shown. B) the surface response plot of the effects of orientation and trac width on stiffness. C) the surface response plot of the effects of trace width and PPXC thickness on stiffness. Least squares model and graphs were generated with JMP Pro software.

Our data suggests that surgical placement of these Neuralace devices should be performed at 90° with respect to the main conformation axis (i.e., the long axis of the target gyrus), as this orientation significantly reduces the stiffness of the device, and thus the forces exerted by it on to brain tissue. Nonetheless, during implantation, surgeons may prefer to handle the structures at the stiffer 0° orientation to avoid damaging the device or to accommodate the physical constraints of the surgery.

### Optimizing the mechanical properties of subdural implants

This study has shown experimentally how the different design factors affect the mechanical properties of these Neuralace devices, and how forces exerted on the brain can be used to evaluate the mechanical performance of subdural implants. The results show that changes in the design factors do impact the mechanical properties of these Neuralace devices in a statistically significant manner. However, it is important to highlight that we are navigating a narrow design parameter space, and the starting point is a design that considers a wider range of constraints such as manufacturing reliability, electrical properties, signal source constraints, and biocompatibility of the material amongst others. For example, the mesh-like open architecture of the Neuralace takes input from a variety of studies which show that such design offers biocompatibility. For subdural implants *Schendel et al*. showed that the distribution of FE depends on the device’s footprint and suggested that a mesh-like open architecture design could reduce the impact of FE on recording performance^13^. Their results indicate that FE on open architectures forms predominately on device-dura interface, rather than on the device-brain tissue interface. Reducing the FE on the device-brain tissue interface is desirable, as it acts as an electrical insulator hindering the recording performance of the electrodes. Navigating the design space while satisfying the technological, physiological, and physical constraints is an active area of neuroengineering research, in which boundaries are constantly pushed with innovative designs, novel materials, and manufacturing techniques. This study portrays one such endeavors and elucidates an experimental approach that can be used in the optimization of conformable neural implant designs with a focus on the forces exerted by the devices on the neural tissue.

## Conclusion

In conclusion, this study highlights the importance of using a DOE framework to systematically investigate the mechanical properties of conformable brain implants, ensuring statistically significant and translatable results. Effects screening can guide the optimization of subdural implant designs in a mechanically aware manner, which can impact the long-term biocompatibility and functionality of the devices. As more devices are designed and implanted, it is crucial to rigorously characterize and report their mechanical properties, enabling broader comparisons between conformable neural implants. This could help clarify the effect of stiffness in the elicited foreign body responses (FBRs) and accelerate the development of long-term conformable neural interfaces.

## Supporting information

Supplement Sections (S1 and S2)

## Acknowledgments

Bethany Miller is thankfully acknowledged for performing ~50 % of the mechanical trials used in this study. Colleen Chemerka is also thankfully acknowledged for manually annotating images pertaining to the structure alignment variability study.

## Conflicts of Interest

Florian Solzbacher declares financial interest in Blackrock Neurotech and Sentiomed, Inc. This potentially competing interest is overseen by the University of Utah’s Conflict of Interest Management.

