## Supplement Sections (S1 and S2) for "Neuralace: Manufacture, Parylene-C Coating, and Mechanical Properties"

### **Supplementary Material**

### S1. Structure alignment reproducibility

Ensuring that the structure is properly aligned with the loading geometries is critical as it impacts the results of the bending experiments. This is due to the squared geometry of the structure as changing the orientation of the structure, with respect to the loading noses, drastically changes the area under strain (i.e., area between the loading noses). Since structure placement was performed manually, and as orientation is one of the controlled experimental factors, it was imperative to characterize the reproducibility of this experimental procedure in order to analyze its effect on the measured mechanical properties. To characterize the reproducibility, two operators performed 10 structure placements on each orientation (0° and 90°) and images were taken from a top-view after structure alignment was performed. During image acquisition a fixed reference frame was added to the images. The obtained images were then manually annotated to extract the coordinates of the fixed reference frame and the structure corners. With these coordinates, the centroid of the irregular polygon defined by the four coordinates was calculated and used as the reference frame to calculate the angle between the x-axis and the top-right corner of the structure (θ). For each structure placement image, the reproducibility was defined as the absolute difference between the average $\bar{\theta}$ and the measured angle θ. This resulted in a mean absolute difference of 1.8° ± 0.58°. This evaluation suggests that the orientation angle can be controlled with a resolution of 3°, considering two standard deviations.

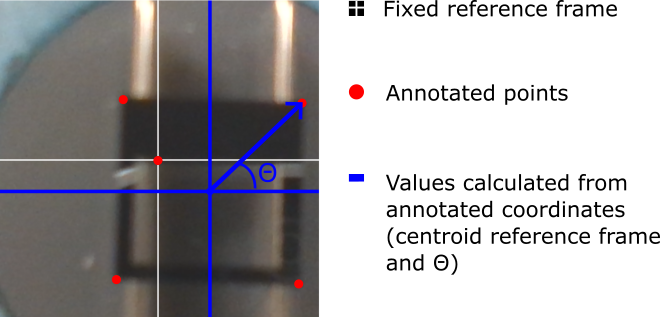

Figure 1: Top view of a Neuralace structure supported by the lower bending geometry, overlaid with manual annotations and calculated reference frame to evaluate the orientation of the structure. Photos were taken from 10 sample placements for each operator to quantify of the orientation placement variation.

#### S2. PPXC Coating

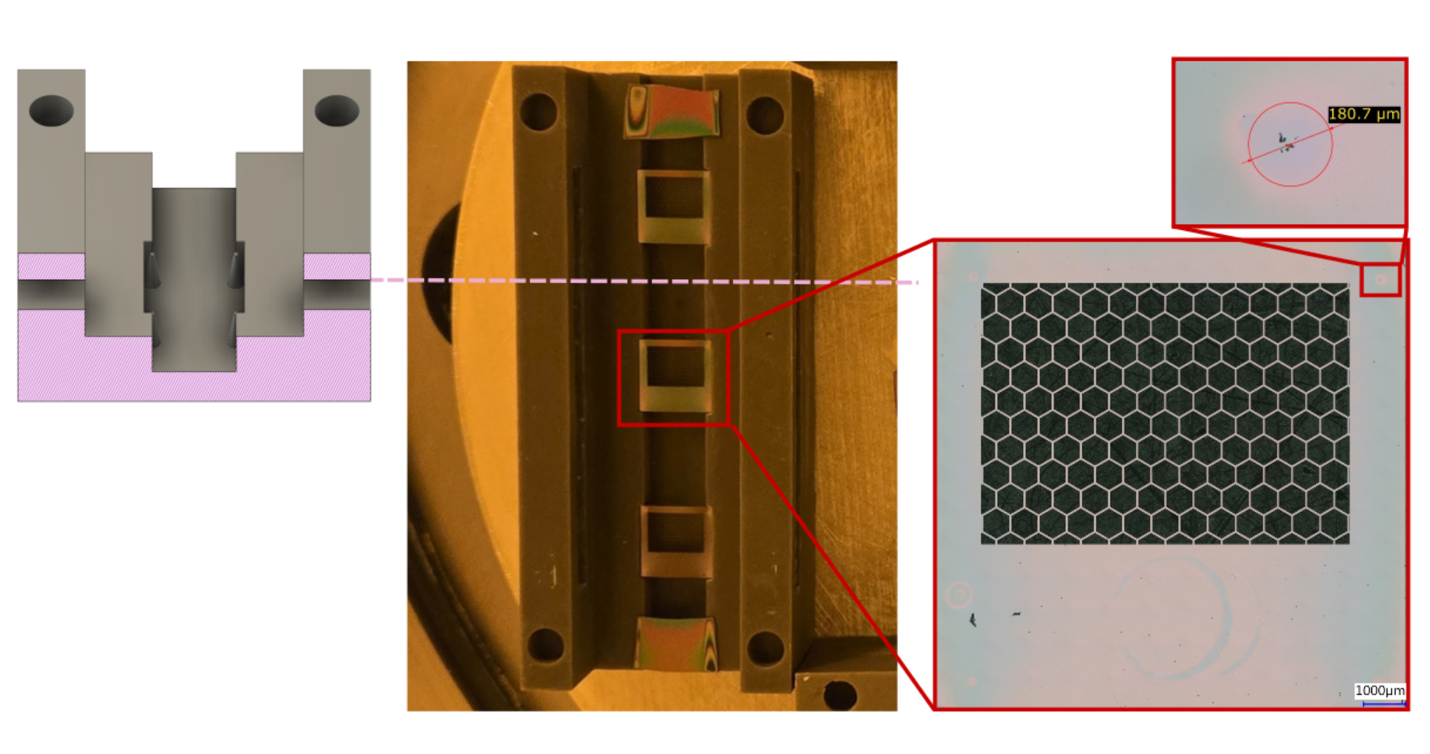

Figure 2: Overview of the parylene-C coating holder. A custom 3D printed holder suspends the lace structures on four prongs. The points of contact in the corner are imaged and measured. After the first coating, the structures are flipped and coated again to ensure complete encapsulation.

To conformally coat the devices with PPXC, structures were suspended on four contact points in custom 3D holder that enables the flow of vaporized dimer above and below the structure. The design of the holder minimizes non-uniformity by minimizing contact area with the structures. Imaging of the coated structures showed that the area affected due to each contact point was 216 ± 69 µm (demonstrative measurement shown in Figure 2). All 12 structures were coated in the same run for consistency, with 8 Si witness chips (WaferPro, CA, USA) beside the structures, two on each structure holder, to help quantify the deposition (see figure 9). Prior to coating, the structures were placed in their holders and plasma cleaned for 2 minutes with an RF setpoint of 1 KW and an Oxygen flow rate of 99 SCCM (March PX-1000, Nordson, OH, USA). After cleaning, the structures were placed on a desiccator and silanized for 2 hours with A-174 Silane, as an adhesion promotion step. The coating process consisted of two consecutive Parylene deposition runs (PDS 2010 Labcoter 2, Specialty Coating Systems, IN, USA), each consuming 1.15 g of solid PPXC precursor dichloro [2,2] paracyclophane. The structures were flipped in between runs to ensure complete encapsulation. The pump-down pressure set point was 10 mTorr during pump down and 25 mTorr during deposition. The vaporizer was set to 175 °C to facilitate sublimation of the precursor and the furnace temperature was set to 690 °C to carry out the pyrolysis of the vaporized precursor. After deposition, the thickness of the PPXC layer on the witness chips was characterized with a profilometer (Dektak Xt stylus, Bruker) to have in an average thickness of 1.78 µm ± 20 nm.
